# Arabidopsis RETINOBLASTOMA-RELATED controls cell size during plant development in a dose-dependent manner

**DOI:** 10.64898/2026.03.30.715244

**Authors:** Rasik Shiekh Bin Hamid, Fruzsina Vadai-Nagy, Magdolna Gombos, Ildikó Domonkos, José Manuel Pérez Pérez, Attila Fehér, Zoltán Magyar

## Abstract

The RETINOBLASTOMA-RELATED (RBR) protein in plants functions as a cell-cycle inhibitor, regulating cell numbers in developing organs and establishing cellular quiescence during growth. Although the role of RBR counterparts in animals also involves regulating cell size, this potential function remains unexplored in plants. We investigated transgenic Arabidopsis plants with altered RBR levels and observed corresponding changes in cell size from embryogenesis through organ development. In addition, stomatal meristemoid cells with reduced RBR levels divided beyond the size threshold, whereas elevated RBR levels increased their size. RBR stimulated terminal differentiation in the stomatal lineage by inducing *MUTE* and *CYCLIN D5;1* expression, whereas reduced RBR levels maintained asymmetric divisions through high *SPEECHLESS* and *CYCLIN D3;1* expression. Interestingly, the cell proliferation-dependent phosphorylation of RBR at the conserved 911Ser site positively correlated with RBR protein levels in the transgenic lines and aligned with the effect of RBR on cell size. This study discusses the potential link between RBR’s control of cell proliferation and cell size, providing new insights into the coordinated regulation of plant development.

## Introduction

Plant development is controlled by the dynamic interaction between cell proliferation and differentiation (Feher and Magyar, 2015; Sablowski, 2016). Cells capable of proliferation can undergo either symmetric or asymmetric division to supply the growing plant or organ with new cells and to establish different cell types. Although cells vary in size, each plant species maintains a consistent distribution of cell sizes across generations, indicating that cell growth is actively regulated during development.

Cell size is closely linked to cellular function, affecting the cell’s physiological, signalling, and mechanical properties of cells (Ginzberg et al., 2015). During cellular proliferation, cell size is determined by the balance between cell growth and division, with feedback mechanisms between these processes maintaining uniformity in cell size. A novel proposed molecular mechanism by which cell size and growth regulate cell cycle transitions involves the dilution of specific cell cycle inhibitors. In budding yeast, cell growth during the G1 phase dilutes the cell cycle inhibitor Whiskey 5 (Whi5), thereby triggering cell cycle entry (Schmoller et al., 2015). In mammals, the functional orthologue of Whi5 is the Retinoblastoma (RB) protein, which is recognised as the first tumour suppressor, and plays a crucial role in regulating cell proliferation, maintaining stem cell status, and controlling cell survival (Du and Pogorlier, 2006; Henley and Dick, 2012). Additionally, RB has been shown to regulate cell size during cell division in a dose-dependent manner, as cell growth dilutes RB during the G1 phase, leading to the onset of the cell cycle (Zatulovskiy et al., 2020). In plants, the RETINOBLASTOMA-RELATED (RBR) proteins have a conserved cell-cycle-inhibitory function (Desvoyes and Gutierrez, 2020). In Arabidopsis, RBR is encoded by a single gene, and its mutation leads to gametophyte lethality (Ebel et al., 2004). The primary effectors of RBR are E2F transcription factors, with three E2Fs (E2FA, E2FB, and E2FC) in Arabidopsis demonstrating RBR-binding capability (Magyar et al., 2005; Magyar et al., 2012; Henriques et al., 2010; Lang et al., 2021; Gombos et al., 2023). Indeed, all three E2Fs trongly associate with RBR, and these E2F-RBR complexes bind to overlapping targets (Lang et al., 2021; Gombos et al., 2023). In the triple *e2fabc* mutant plant, the RBR was notably unable to establish and maintain stable cellular quiescence, leading to the development of enlarged organs composed of a greater number of cells that are comparable in size to the control WT (Gombos et al., 2023). Plant RBR is mainly found in the nucleus and controls the expression of genes involved in both the G1-S and G2-M phases of the cell cycle. Cell size regulation in plants is considered to occur during these cell cycle transitions (Jones et al., 2016). Recent research has indicated that CDK-inhibitory proteins, specifically KIP-RELATED PROTEINS (KRPs) and SIAMESE (SIM) and SIAMESE-RELATED (SMR) proteins, operate in distinct G-phases, G1 and G2, respectively. Notably, chromatin-bound KRP4 regulates the transition from the G1 to the S phase in a concentration-dependent manner, suggesting that its dilution in proliferating G1 cells determines cell size at the time of cell cycle entry (D’Ario et al., 2021). In a separate molecular mechanism, the MYB3R-4 transcription factor induces the expression of the GRASS transcription factor SCARECROW-LIKE 28 (SCL28), which, together with the APETALA 2 (AP2)-type protein SMALL ORGAN SIZE 1 (SMOS1), promotes the expression of SMR proteins (Nomoto et al., 2022). The upregulated SMRs inhibit entry into the M-phase by modulating G2-M phase-specific CDKs, thereby shifting the cell cycle to the endocycle and thus controlling cell size. Interestingly, the F-BOX-LIKE 17 (FBL17), which promotes the degradation of KRPs, and the activator MYB3R-4 are targets of the RBR-E2F pathway (Zhao et al., 2012; Desvoyes & Gutierrez, 2020; Gombos et al., 2023). Furthermore, RBR activity is believed to be regulated by CDKs, as the phosphorylation of RBR hinders its interaction with E2Fs (Magyar et al., 2012; Pettkó-Szandtner et al., 2026). Recent research has shown that plant RBR is phosphorylated at multiple CDK sites, suggesting that different CDKs may phosphorylate RBR both in complexes with E2Fs and in monomeric forms (Pettkó-Szandtner et al., 2026). The S6 kinase (S6K), regulated by the TARGET OF RAPAMYCIN (TOR) kinase, has also been suggested to function as an RBR kinase (Henriques et al., 2010; 2013). This pathway is involved in regulating cell size in response to nutrient availability (Ahmad et al., 2019).

These data collectively suggest that RBR may regulate cell size in plants; however, its role as a cell sizer remains to be examined. In the green algae Chlamydomonas, the RBR homologue MAT3 has been identified as a size-dependent repressor, ensuring that cells reach a minimum size before division. The absence of MAT3 results in a phenotype characterised by smaller cells (Umen and Goodenough, 2001). Notably, a substantial reduction in RBR levels during the early phases of plant development leads to excessive proliferation of stem and progenitor cells, thereby inhibiting differentiation and inducing spontaneous cell death (Borghi et al., 2010; Gutzat et al., 2011; Gombos et al., 2023). Conversely, its overexpression has the opposite effect, completely inhibiting proliferation and causing the loss of stem cell activity (Wildwater et al., 2005; Wyrzykowska et al., 2006). Therefore, the sensitivity of plant growth and development to variations in RBR levels poses challenges in determining RBR’s various functions in plant development, including cell size regulation.

In this study, we examined the role of RBR in regulating cell size during Arabidopsis development using two transgenic lines with altered RBR levels, without inducing macroscopic changes during sporophyte development. Changes in cell size in the meristem and developing organs followed changes in RBR levels; which increased in the ectopic RBR-expressing line or decreased in the *rbr1-2* mutant. Furthermore, stomatal meristemoid cells are sensitive to RBR levels; in the *rbr1-2* mutant, these cells divide at a smaller size, thereby surpassing the cell-size threshold observed in the WT control. In the ectopic RBR-expressing line, these cells undergo fewer divisions and are larger. Ectopic RBR stimulates terminal differentiation within the stomatal lineage by inducing the expression of the *MUTE* transcription factor and its downstream target, *CYCLIN D5;1* (*CYCD5;1*), implicated in the transition from meristemoid to guard mother cell. Conversely, reduced RBR levels sustained asymmetric divisions in the stomatal meristemoid cells through elevated *SPEECHLESS* (*SPCH*) and *CYCLIN D3;1* (*CYCD3;1*) expression. These data indicate that RBR determines the proliferation activity and the size of meristemoid cells and may regulate cell fate within the stomatal lineage. Notably, inhibitory phosphorylation of RBR followed its level in these transgenic lines, being lowest in the *rbr1-2* mutant and the highest in the ectopic line. These data support the idea that dose-dependent phosphorylation may affect the timing of cell cycle entry and cell size.

## Material and methods

### Plant material and growth conditions

Wild-type (WT) *Arabidopsis thaliana* ecotype Columbia-0 (Col-0) along with transgenic T-DNA insertion *rbr1-2* mutant and ectopic RBR-GFP expressing seeds (Hamid et al., 2025), were sterilized using commercial bleach. These seeds were then re-suspended in sterile water, and subjected to cold treatment at 4 °C in the dark for two days (Clough and Bent, 1998). Unless otherwise specified, the plants were cultivated under continuous light at 22 °C *in vitro* on half-strength germination medium, with a light intensity of 100 μmol mE^-2^ s^-1^. The first leaf pairs of these lines were cultivated *in vitro*, harvested between 7 and 16 days after germination (DAG), flash frozen, and stored at -80 °C.

### Immunoblotting assays

Immunoblotting assays were performed as described previously (Gombos et al., 2023). The primary antibodies used in these assays were anti-phospho-specific RB (Ser^807/811^) rabbit polyclonal antibody (Cell Signalling Tech), and anti-Arabidopsis N-terminal specific RBR mouse polyclonal antibody (Hamid et al., 2025), anti-Arabidopsis C-terminal specific RBR chicken polyclonal antibody (Agrisera).

### Confocal laser and scanning electron microscopy (SEM) observations

To analyse root samples using confocal laser microscopy (SP5, Leica), the seedlings were cultivated on vertically oriented plates, and the roots were stained with propidium iodide (PI; 20 μg/mL) before being photographed. Cell length was measured using the Image J software. Cotyledon and leaf samples were similarly stained with PI, and the epidermis was imaged using confocal microscopy. Mature dried seeds were dissected, and the cotyledon epidermis was stained with PI and photographed as previously described (Leviczky et al., 2019).

Young seedlings were fixed in a 9:1 ethanol to acetic acid solution and cleared with Hoyer’s solution (a mixture of 100 g chloral hydrate, 10 g of glycerol, 15 g of gum arabic, and 25 mL of water). Microscopic observations were performed using the first leaf. After whole leaf images were captured, palisade cells were observed using a differential interference contrast (DIC) microscope (Leica SP5). The captured images were analysed using the Image J software (ver.2.1.0; rsb.info.nih.gov/ij) and the average size of palisade cells and the number of cells in the uppermost layer of palisade tissue per leaf, were calculated according to methods previously described methods (Nomoto et al., 2022).

Young seedlings were vacuum infiltrated and fixed with 100% methanol for 20 min, dehydrated in 100% ethanol for 30 min and then in fresh 100% ethanol overnight. The following day, the samples were critical point dried, mounted on SEM stubs, and observed in a JEOL JSM-7100F/LV scanning electron microscope in the low-vacuum mode. High-contrast cell outlines in uncoated plants were imaged according to Talbot and White (2013) by detecting backscattered electrons at 15 kV 41 accelerating voltage and 40 Pa pressure in the specimen chamber. Cell size was measured using the Image J software.

### Quantitative RT-PCR

RNA was extracted from the first leaf pairs of Arabidopsis, harvested at various developmental stages, specifically at nine or twelve days post-germination using the CTAB-LiCl method as described by Jaakola et al. (2001). RNA samples were treated with DNase1 (ThermoScientific #EN0521) in accordance with the manufacturer’s protocol. cDNA was synthesized from 1 μg of RNA using the ThermoScientific Reverse Transcription Kit (#K1691) with random hexamers, following the manufacturer’s guidelines. A control reaction devoid of the RevertAid enzyme was conducted to ensure the absence of genomic DNA contamination. RT-qPCR was performed using SYBR Green (TaKaRa TB Green Primer Ex TaqII, #RR820Q) as per the manufacturer’s instructions, on a BioRAD CFX 384 Thermal Cycler (BioRAD) with the following parameters: 50 °C for 2 min, 95 °C for 10 min, 95 °C for 15 s, 60 °C for 1 min, 40 cycles, followed by melting point analysis. Each reaction was conducted in triplicate, and specificity was confirmed by a single peak in the melting curve. Data normalization was performed using the average Ct value of two housekeeping genes (ACTIN2 and UBIQUITIN CONJUGATING ENZYME 18). The specific primer sequences used are detailed in Supplementary Table 1.

## Results

### Altering RBR levels influences the average cell size in differentiated organs

To clarify the role of RBR as a determinant of cell size, we examined its effects on cell size in organs with determinate growth in two transgenic lines in which RBR levels were either elevated (ectopic RBR-GFP line driven by its own promoter) or reduced (*rbr1-2* T-DNA insertion mutant line) (Hamid et al., 2025), and compared them with the WT control. Our study focused on the first leaf pair, measuring the size and number of palisade cells in *in vitro*-grown WT, RBR-GFP, and *rbr1-2* lines during their development from 7 to 14 days after germination (DAG). We also monitored the fluorescence signal of RBR-GFP, which was ubiquitously distributed throughout the palisade cells and concentrated in the nuclei (Fig. 1A). At all developmental stages examined, the size of palisade cells in the first leaf correlated with RBR levels. Compared with the control wild type (WT), the cells in the ectopic RBR-GFP line were significantly larger, whereas those in the *rbr1-2* line with reduced RBR level were notably smaller (Fig. 1B-D). The smallest category of palisade cells (50-100 µm^2^) from the first leaf at 7 DAG was observed exclusively in the *rbr1-2* mutant, whereas the largest cells (400-500 µm^2^) were found only in the ectopic RBR-GFP line (Fig. 1E). The results indicate that cell number correlated with cell size depending on the level of RBR: more but smaller cells per area characterise the *rbr1-2* mutant, whereas fewer but larger cells are found in the RBR-GFP line, compared to the control. To further confirm that RBR influences cell size in organs with determinate growth, we examined similar parameters in the petals of RBR transgenic lines and compared them with those of the WT control (Suppl. Fig. 1A). The size of palisade cells in petals, consistent with leaf data, displayed a strong correlation with RBR levels (Suppl. Fig 1B). As expected, cell numbers were inversely related to RBR levels, whereas cell size was not (Suppl. Fig. 1C).

**Figure 1.**
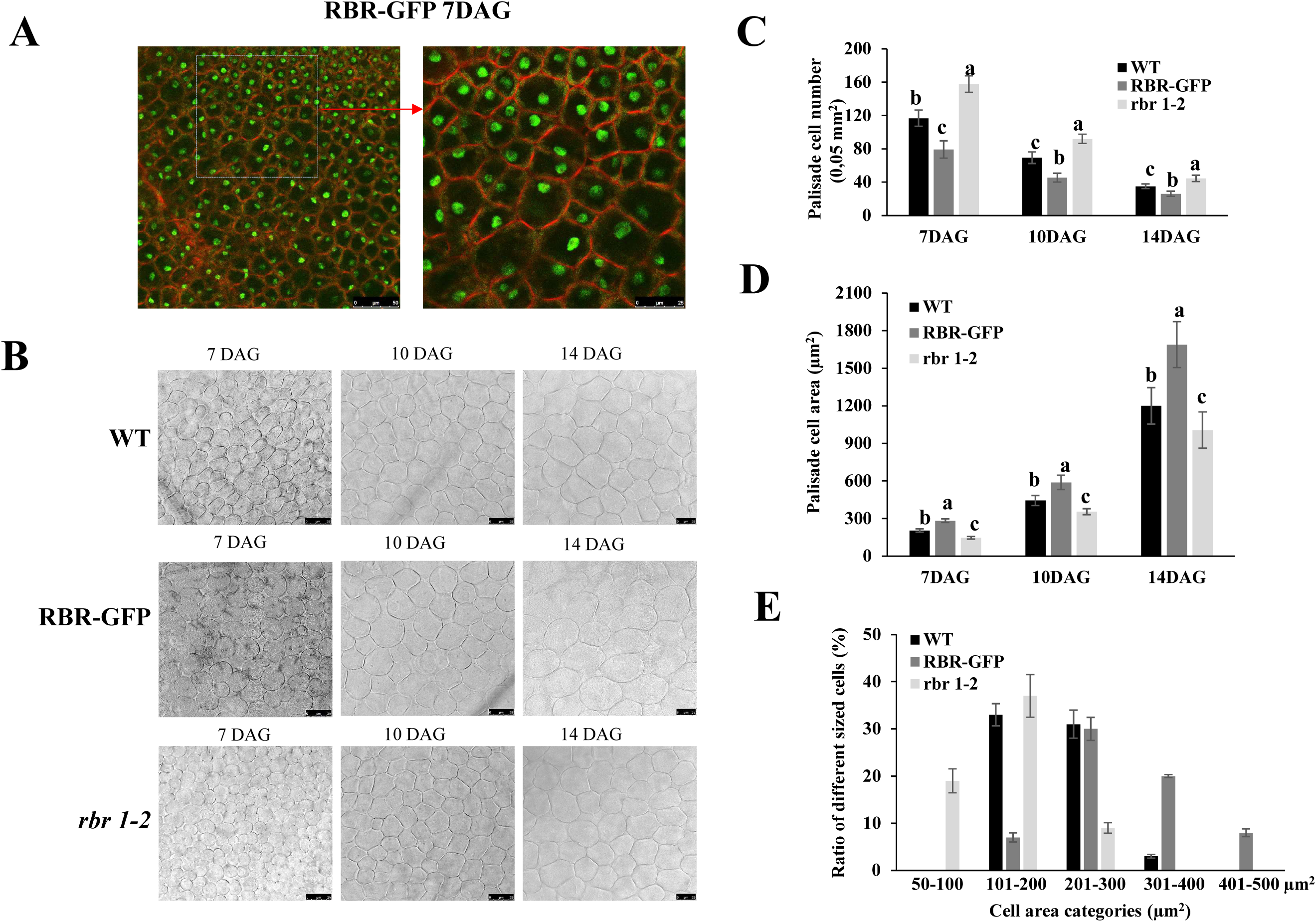
The concentration of RBR affects cell size in organs characterised by determinate growth. **A,** The first leaf pair of the RBR-GFP line was stained with propidium iodide and imaged using confocal laser microscopy. The scale bars are 50 and 25 µm, respectively. **B,** Images of palisade cells of WT, RBR-GFP and *rbr1-2* first leaf pairs at 7, 10 and 14 days after germination (DAG). Scale bar is shown (25 µm). **C,** The number of cells is 0,05 mm2 squares of the first leaves (n=5) for the investigated lines. **D,** The average area of palisade cells was determined from five leaves per line, each leaf with over 40 cells analysed. **E,** The ratio of different-sized cells at 7 DAG was plotted. Different letters indicate statistically significant differences (p < 0.05) according to on one-way ANOVA with Tukey’s HSD post-hoc test.

In conclusion, when the RBR level is altered in Arabidopsis organs, the size of differentiated cells varies accordingly. These size differences correlate with cell proliferation but are maintained during the post-mitotic developmental phase, suggesting that RBR may regulate the growth of differentiated cells.

### RBR level influences cell size, cell division, and differentiation within the stomatal lineage

To further investigate the regulatory effect of RBR levels on cell size, we assessed their impact on dividing plant cells characterised by specific cell sizes. In the epidermis of cotyledons and true leaves, triangular-shaped stomatal meristemoid cells undergo asymmetric divisions before switching to symmetric division and differentiating into guard cells (Kim and Torii, 2023). These meristemoids gradually decrease in size due to further asymmetric divisions until they reach a critical threshold that triggers symmetric division (Gong et al., 2023; Kim and Torii, 2023). We hypothesised that RBR would alter the cell size threshold in a dose-dependent manner. Previous studies have confirmed the regulatory role of RBR within this lineage (Borghi et al., 2010; Yang et al., 2014; Kim and Torii, 2023). Using RBR-GFP as a reporter, we confirmed the presence of RBR-GFP in all epidermal cells of the cotyledon and the first leaf, including those of the stomatal lineage (Suppl. Fig. 2; Matos et al., 2014; Őszi et al., 2020). We analysed the leaves and cotyledons in WT, RBR-GFP, and *rbr1-2* lines using confocal microscopy after propidium iodide (PI) staining (Fig. 2A-C and Suppl. Fig. 3A-D). First, an analysis of epidermal cell size distribution across different categories was conducted for all three lines. Compared with the other two lines, the *rbr1-2* line presented the smallest epidermal cells, whereas a notable shift from smaller to larger cells was observed in the ectopic RBR line (Fig. 2D and Suppl. Fig. 3E). Attention was then focused on triangular-shaped meristemoid cells. These cells were smallest in the *rbr1-2* line (Fig. 2E; Suppl. Fig. 3F), representing a new cell-size category (Fig. 2D; Suppl. Fig. 3E). In contrast, these meristemoid cells were larger in the RBR-GFP line than in the WT control. The *rbr1-2* line showed notably smaller, more numerous cells than the WT control, indicating extra asymmetric divisions at reduced cell size (Fig. 2E and F; Suppl. Fig. 3F and G). Conversely, in the ectopic RBR line, cell size shifted toward the larger category compared to the WT, indicating fewer cell divisions and resulting in larger cells relative to the control. Notably, the expression of the cell cycle marker genes *ORIGIN RECOGNITION COMPLEX, SUBUNIT 2* (*ORC2*), specific to the S-phase, and *CYCLIN-DEPENDENT KINASE B1;1* (*CDKB1;1*), relevant to the G2-M-phase, correlated with the observed RBR-dependent differences in cell proliferation in the first leaves (Fig. 2G, H).

**Figure 2.**
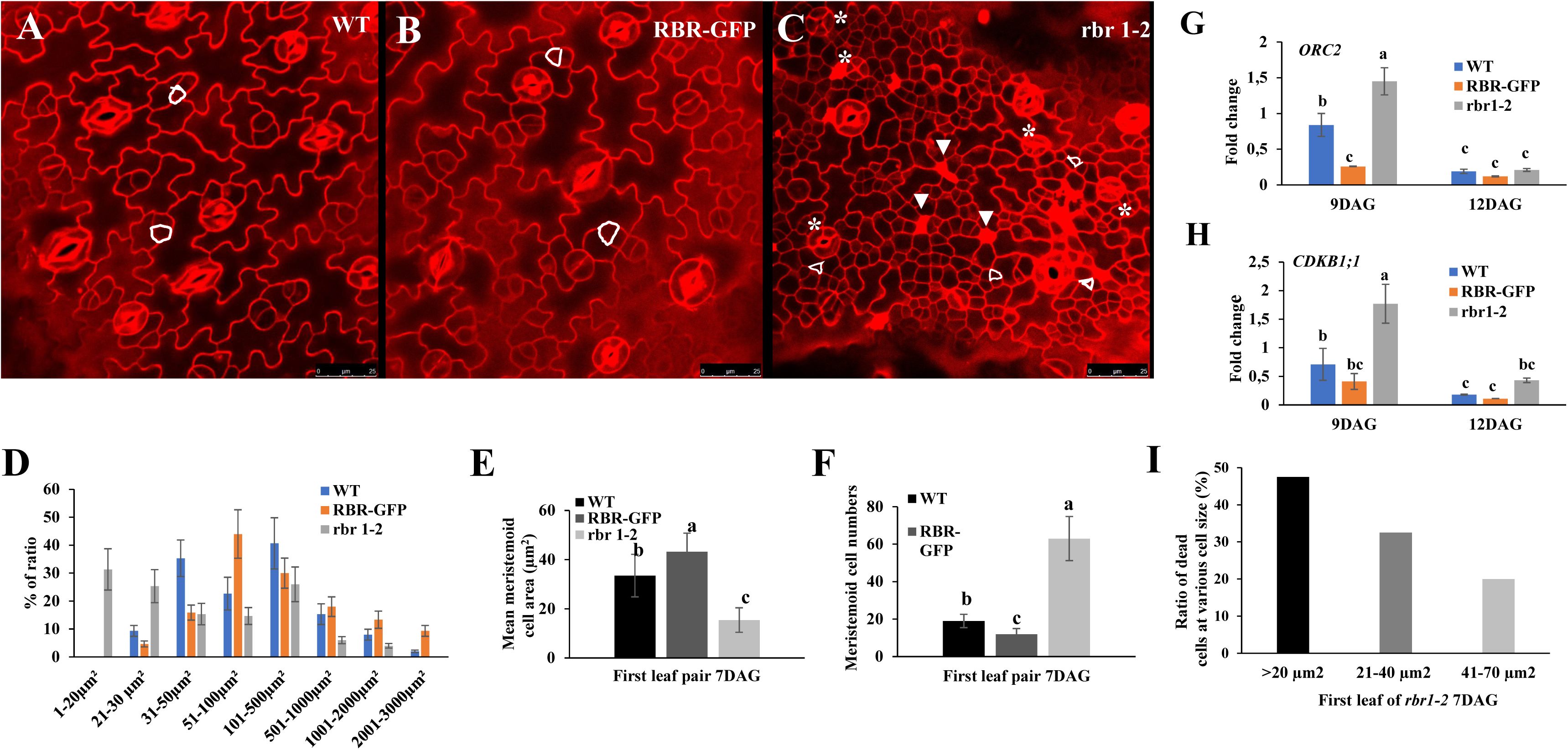
RBR dependent control of cell size and number in developing leaves. **A-C,** Representative confocal laser microscopy images of propidium iodide-stained first leaf pairs of the WT (**A**), RBR-GFP (**B**) and *rbr1-2* (**C**) lines at 7 days after germination (DAG) are presented. The triangular cells with white contours are meristemoid-like cells. Asterisks denote stomata with more than two guard cells, while arrowheads indicate few dead cells in the *rbr1-2* line. The scale bars represent 25 μm. D, Ratio of cells in different size categories across the whole leaf (150 cells counted per image) in all three lines. E, The mean cell area of the stomatal meristemoids at 7 DAG was calculated and plotted. F, Meristemoid cell number was also counted and plotted in all three lines. The values are averages from ten areas analysed (n=10). **G-H**, The regulation of *ORIGIN RECOGNITION COMPLEX, SUBUNIT 2* (*ORC2*), which is specific to the S-phase (**G**) and the *CYCLIN-DEPENDENT KINASE B1;1* (*CDKB1;1*) pertinent to the G2-M-phase (**H**), is influenced by the levels of RBR in agreement with their effect on cell proliferation (values represent fold changes normalised to the value of the relevant transcript in seedlings at 9 DAG, which was arbitrarily set to 1. The data are the means ± SD. N=3 biological replicates). Different letters indicate statistically significant differences (p < 0.05) according to on one-way ANOVA with Tukey’s HSD post-hoc test. **I**, The size of the dead cells in the epidermis of *rbr1-2* mutant leaves was determined (n=40 dead cells/leaf), and the ratios of the three size categories were calculated and plotted.

Consistent with previous findings on the role of RBR in cell survival (Borgi et al., 2010; Gombos et al., 2023), numerous dead small meristemoid-like cells were observed after PI staining of epidermal cells, particularly in the *rbr1-2* mutant (Fig. 2C and Suppl. Fig. 3D). Although the smallest meristemoid cells may have a higher rate of cell death, no correlation was observed between cell death and cell size (Fig. 2I). Furthermore, in the epidermis of the *rbr1-2* leaf, differentiated guard cells consisting of three or four cells, rather than the usual two, were observed (Fig. 2C and Suppl. Fig. 3C, D). This finding supports that RBR is required for the establishment and maintenance of quiescence in these differentiated cells (Gombos et al., 2023).

The differential effect of various RBR levels on stomatal meristemoid cells was further confirmed by examining the epidermis of mature embryonic cotyledons (Suppl. Fig. 4), where epidermal cells remain undifferentiated and arrested at different meristemoid-like stages (Kim and Torii; 2023). Overall, the data suggest that the level of RBR influences the size of stomatal meristemoid cells in the epidermis during both embryonic and post-embryonic stages, in cotyledons and true leaves. In line with the increased number of stomatal meristemoids, more differentiation of stomatal guard cells was observed in the leaf epidermis of the *rbr1-2* line compared to the WT control (Suppl. Fig. 5A-D). Conversely, fewer guard cells developed in the ectopic RBR-GFP line relative to the WT (Suppl. Fig. 5D). Interestingly, the size of guard cells was not affected by the higher RBR level; however, they appeared to differentiate with a smaller size in the *rbr1-2* line (Suppl. Fig. 5E).

The progression of the stomatal lineage is controlled by basic helix-loop-helix (bHLH) transcription factors, SPEECHLESS (SPCH), MUTE, and FAMA, along with their dimeric partners (MacAlister et al., 2007; Kim & Torii, 2023). SPCH initiates asymmetric divisions within protodermal cells, leading to the formation of meristemoids that act as self-renewing stomatal precursors. MUTE subsequently terminates this self-renewing activity and guides the cell toward differentiation. We examined the expression of *SPCH* and *MUTE* in the first leaf pair of 9- and 12 DAG (Fig. 3A, B). The expression of *SPCH* was higher in young leaves at 9 DAG in a manner dependent on RBR concentration; increased expression was observed in the *rbr1-2* mutant compared to WT and the ectopic RBR-GFP expressing line (Fig. 3A). In contrast to *SPCH*, the expression of *MUTE* increased most in the leaf of the RBR-GFP line at 9 DAG, whereas it was repressed in the *rbr1-2* compared to the WT (Fig. 3B). Recent research demonstrated that SPCH and MUTE target different G1-type CYCLIN D genes; SPCH activates *CYCLIN D3;1* (*CYCD3;1*), which is involved in the activation of asymmetric divisions, whereas *MUTE* stimulates the expression of *CYCD5;1* and *CYCD7;1* (Kumar et al., 2018; Han et al., 2022). Both D-type cyclins regulate the transition of meristemoid cells to guard mother cells by activating symmetric divisions. These three D-type CYCLINs showed different regulation patterns in the *rbr1-2* and ectopic RBR-GFP lines; *CYCD3;1* was increased in the *rbr1-2* line, whereas *CYCD5;1* and *CYCD7;1* had the most notable rise in the RBR-GFP line compared to the WT control (Fig. 3C-E). These data indicate that RBR not only controls the proliferation activity and size of meristemoid cells but may also affect cell fate within the stomatal lineage. Specifically, higher RBR levels promote the final symmetric division by stimulating the expression of *MUTE*, whereas lower RBR levels sustain asymmetric divisions by maintaining high levels of *SPCH* expression.

**Figure 3.**
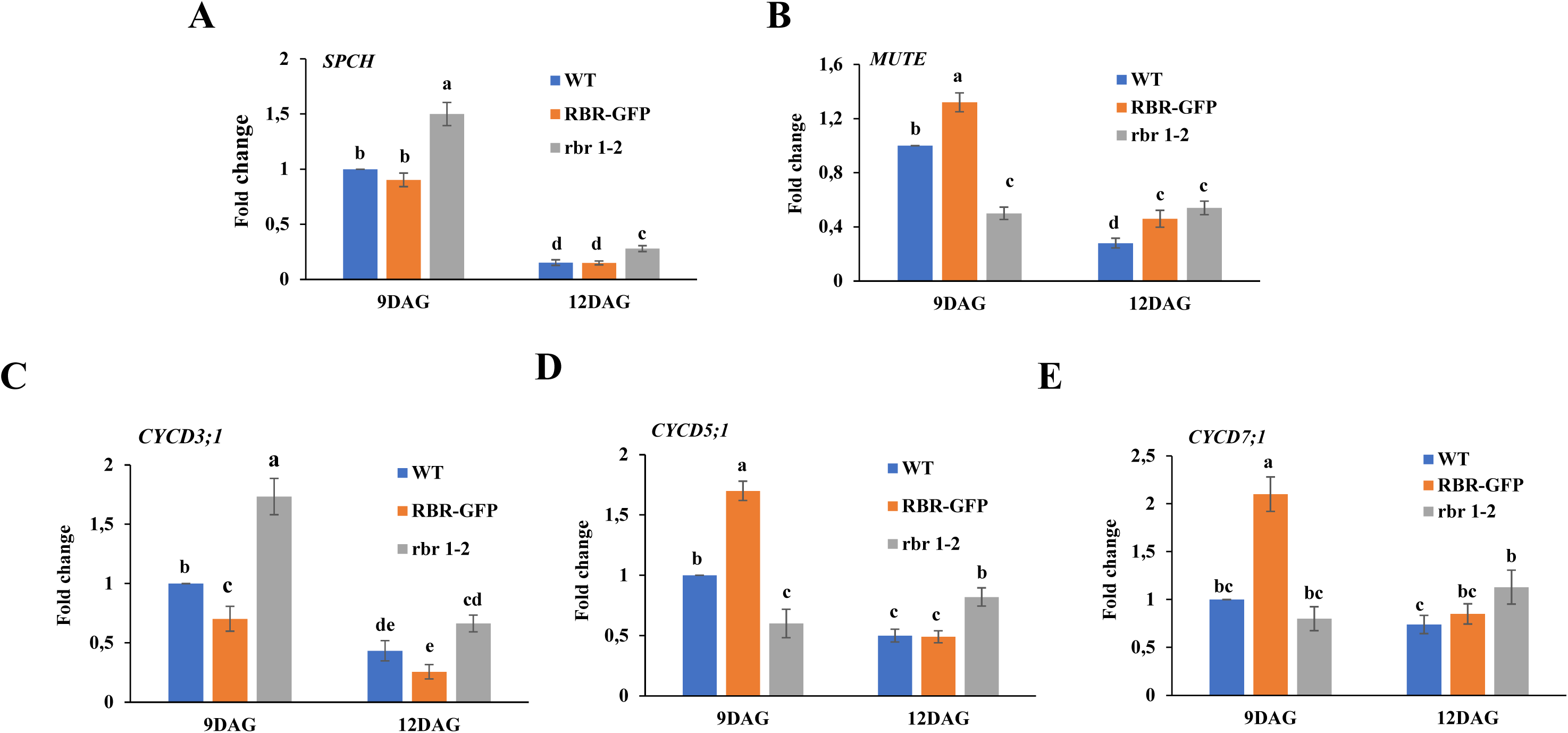
The RBR level influences the expression of genes controlling the division and differentiation of meristemoids. **A-B,** *SPEECHLESS (SPCH)* (**A**) and *MUTE* (**B**) regulation inversely related to the RBR levels in the first leaf pair after 9 days of germination (DAG). The expression of the SPCH-activated CYCD3;1 gene **C**, The expression of the SPCH-acivated *CYCD3;1* gene, which regulates asymmetric cell division, is negatively influenced by RBR levels. **D-E,** Ectopic RBR enhances the expression of the downstream D-type cyclin genes *CYCD5;1* (**D**) and *CYCD7;1* (**E**) of the symmetric cell division and cell differentiation-promoting MUTE transcription factor, which governs the transition from meristemoid to guard mother cells. The values represent the fold changes normalised to the value of the relevant transcript in the seedlings at 9DAG, which was arbitrarily set to 1. The data are the means ± SD. N=3 biological replicates. Different letters indicate statistically significant differences (*p* < 0.05) according to one-way ANOVA with Tukey’s HSD post-hoc test.

### Cells in the root meristem divide at varying cell sizes depending on the level of RBR

To further support the possibility that RBR modulates cell size in dividing cells, we examined its dose-dependent impact on cell size within the root of two RBR transgenic Arabidopsis lines relative to the wild-type (WT) control.

The root is a continuously growing organ of indeterminate size, as its meristem consists of renewable stem cells maintained in a specific environment called the stem cell niche (SCN). Stem cells are attached to quiescent cells (QC), and they divide asymmetrically to produce initials for all the cell layers that make up the root tissues. Their daughter progenitor cells undergo symmetrical division within the proximal root meristem (RM) to supply the expanding root with new cells. Meanwhile, cells even further from the QC stop proliferating and begin to elongate as they reach the transition zone (Baluska et al., 1996). Confirming previous findings, the GFP signal from the RBR-GFP expressing line shows a widespread pattern throughout the root tip and in all cells of the root meristem, including the transition zone, as well as in differentiated columella cells (Fig. 4A, and Magyar et al., 2012; Őszi et al., 2020). In all locations, it predominantly localises in the nuclei. The nuclear RBR-GFP signal was absent only from dividing cells, presumably those in the M-phase, in agreement with previous observations (Fig. 4A, and Miguel-Hernández et al., 2024). Arabidopsis roots of the WT, RBR-GFP, and *rbr1-2* lines were grown together on vertical plates for five days post-germination, during which the root lengths were found to be comparable. To assess whether there were variations in cell size within the root meristems (RM), the root tips were visualised under a confocal laser microscope following PI staining (Fig. 4B). The lengths of cortex cells were measured along the root tip and plotted according to the distance from the QC (Fig. 4C). Cells of the RBR-GFP line began to elongate closer to the QC than those of the wild type, whereas cells of the *rbr1-2* line show the opposite result. Furthermore, the average length of the dividing cortex cells varied with RBR levels; it was observed that, compared to the WT, it increased in the ectopic RBR-GFP, while it decreased in the *rbr1-2* mutant (Fig. 4D). Consistent with this, the number of cortical cells in the RM was inversely related to RBR levels (Fig. 4E), resulting in differences in root meristem size (Fig. 4B).

**Figure 4.**
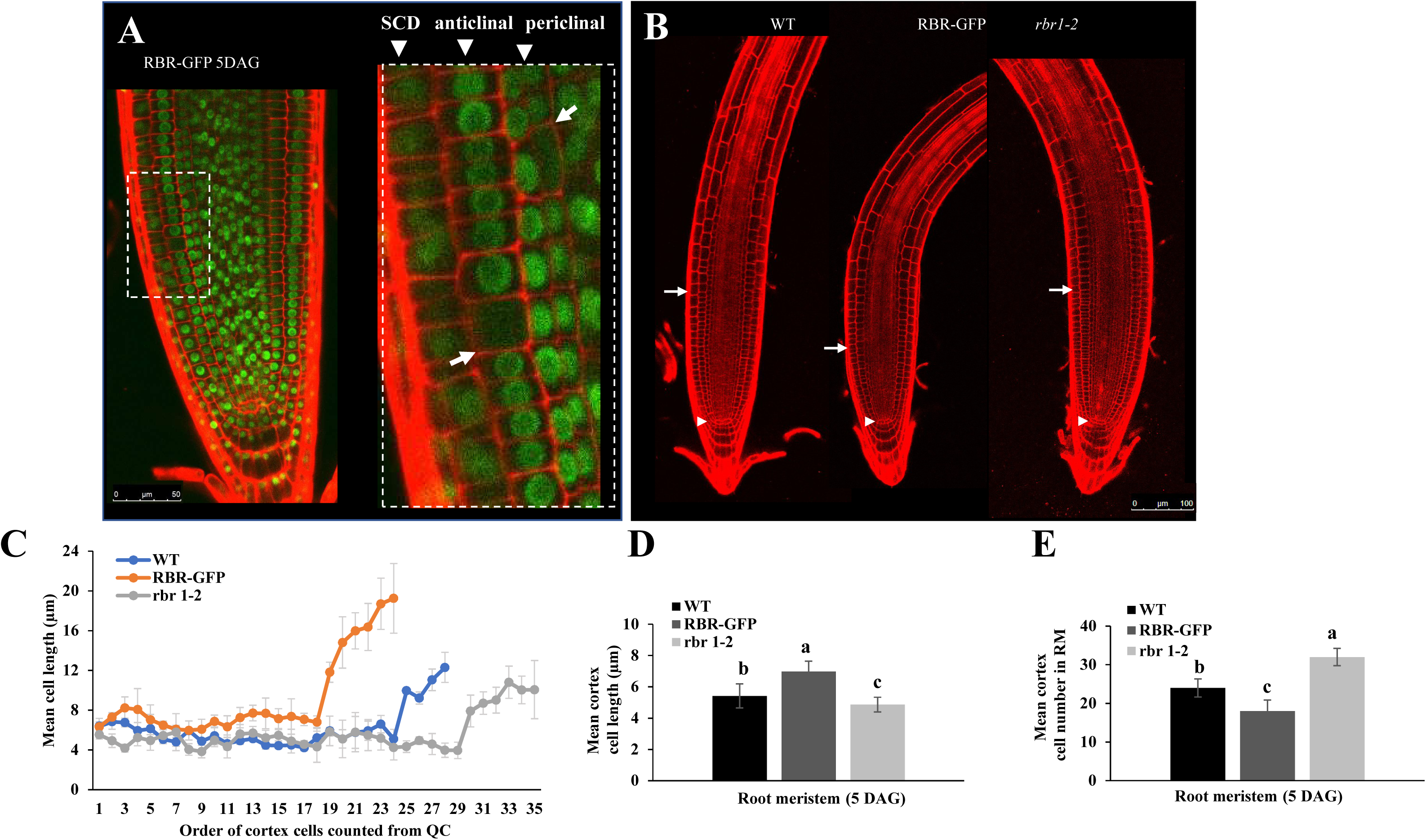
The nuclear RBR protein is widely distributed throughout the root meristem (RM) and influences cell length in a dose-dependent manner. **A**, The GFP signal from RBR-GFP is present throughout the root tip, highlighting all cells in the RM as well as the non-dividing zones. Arrowheads indicate cell files, while arrows point to cells presumably in M-phase that lack nuclear RBR-GFP signal. **B**, Images of propidium iodide-stained roots of WT, RBR-GFP and *rbr1-2* seedlings, grown for 5 DAG, were analysed using confocal microscopy. The scale bar represents 100 µm. Arrowheads point to quiescent centre cells, while arrows mark the end of the dividing zone in the RM. **C**, The spatial distribution of cell length along root meristems is shown for all the lines. **D**, The mean cell length was calculated along the cortical cell file from the quiescent centre to the transition zone. E, Cortex cells in the root meristem were counted, and their number was plotted. For C-E, the values are averages from different plants (n=10, mean±SD). Different letters indicate significant differences (*p* < 0.05) according to ANOVA with Tukey’s HSD test.

These data suggest that the size at which progenitor root cortex cells divide in the meristem depends on RBR levels; they divide at larger or smaller sizes, corresponding to increased or decreased RBR levels, respectively.

### The inhibitory phosphorylation of RBR correlates with its overall level

RBR is regulated through phosphorylation, and phosphorylation at the conserved 911S site is considered to be indicative of an inactive, multi-phosphorylated RBR, which is unable to form a complex with E2F transcription factors (Magyar et al., 2012; Pettkó-Szandtner et al., 2026). Previous research has demonstrated that the phosphorylation of RBR at the 911S site depends on the cell cycle (Ábrahám et al., 2011) and is found in meristems and young, developing organs rich in proliferating cells (Magyar et al., 2012; Őszi et al., 2020; Gombos et al., 2023; Pettkó-Szandtner et al., 2026). Therefore, we chose to investigate how RBR phosphorylation correlates with cell division and cell size. RBR and P-RBR911S levels were evaluated in ectopic RBR-GFP and *rbr1-2* mutant seedlings and compared with the control WT using a western blot assay with RBR and P-RBR-specific antibodies (Fig. 5). The P-RBR911S signal closely matched RBR levels, being most prominent in RBR-GFP and least in *rbr1-2* mutant seedlings compared to the WT (Fig. 5A, B). Given that cell number in developing organs inversely correlates with RBR abundance (Fig. 2, Suppl. Fig. 3; Wyrzykowska et al., 2006; Gutzat et al., 2011; Borghi et al., 2010; Gombos et al., 2023), these findings suggest that an increase in RBR levels is paralleled by a higher phosphorylation rate, which is necessary to inhibit its function as a cell cycle inhibitor. To further support the hypothesis that RBR phosphorylation depends on its level in proliferating cells, we examined RBR and P-RBR levels at various stages of leaf development in the *rbr1-2* mutant and an RBR-GFP-expressing line, compared with WT, using a western blot assay. A sharp transition from cell proliferation to expansion occurs in Arabidopsis leaves between days 10 and 13 DAG (Andriankaja et al. 2012; Fig. 2G, H). The patterns of RBR levels were consistent across the three lines, with the highest levels observed during the proliferative developmental stage of young leaves at 9 DAG, followed by a decline as cell-division activity decreases in the leaves (Fig. 5E, F). The levels of P-RBR closely corresponded to those of RBR, with the highest P-RBR levels observed in the RBR-GFP line and the lowest in the *rbr1-2* mutant, relative to the WT control (Fig. 5E, F). Consequently, the P-RBR level was indicative of leaf cell proliferation activity within each investigated line, regardless of the overall RBR level. However, it was inversely correlated with differences in cell proliferation among the investigated lines (Fig. 2F, H; Fig. 5E, F). Altogether, our findings suggest that phosphorylation of RBR at 911S is correlated with overall RBR levels in seedlings and developing leaves.

**Figure 5.**
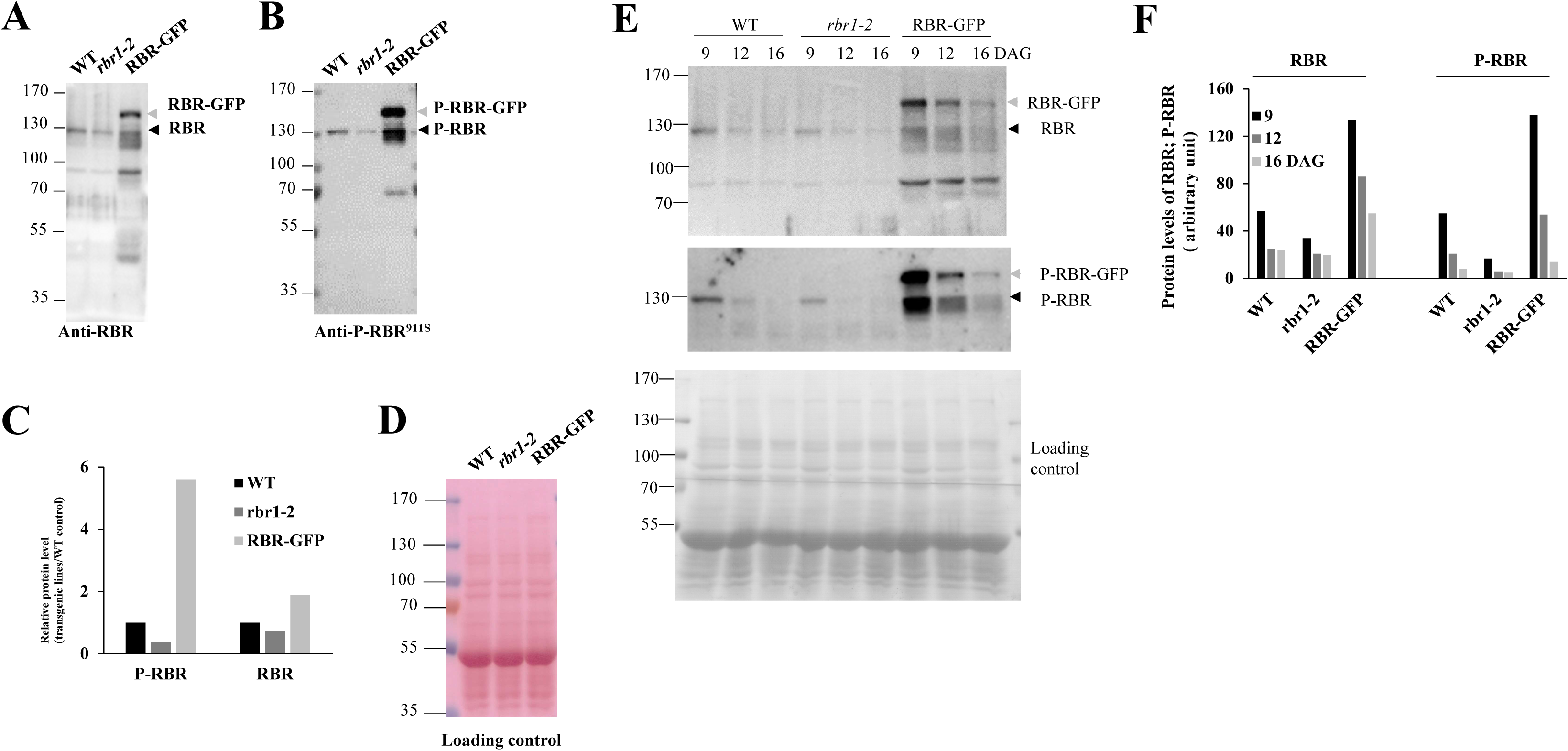
The phosphorylation level of RBR at the 911S site correlates with RBR protein abundance. **A-D**, Western blot shows RBR (**A**) and P-RBR^911S^ (**B**) protein levels in seven-day-old seedlings of WT, *rbr1-2*, and RBR-GFP lines. The band intensities were measured and are shown as relative protein levels (**C**). Ponceau-stained membrane is shown to confirm equal loading of total proteins (**D**). **E-F**, An immunoblot assay was conducted on the first leaf pairs purified from WT, rbr1-2, and RBR-GFP lines at 9, 12, and 16 days after germination (DAG), utilising RBR and P-RBR antibodies (**E**). The relative protein levels of RBR and P-RBR are shown as corresponding band intensities (**F**). Molecular weight markers are indicated on the right side of the blots. The grey arrowheads denote GFP-fused RBR and P-RBR, whereas black arrowheads indicate endogenous RBR and P-RBR levels.

## Discussion

Based on earlier experimental observations, we hypothesized that cell size in Arabidopsis plants could be influenced by RBR protein levels. The data presented here show that the size of dividing cells in the root meristem, embryonic and post-embryonic cotyledons, and true leaves is indeed influenced by RBR levels. Increased RBR in ectopic RBR-GFP expressing seedlings enlarges dividing-cell size, whereas decreased RBR in *rbr1-2* mutant plants reduces the size threshold for division.

We demonstrate that cells in the proximal Arabidopsis RM divide at different sizes depending on whether RBR levels are increased or decreased in RBR-GFP and *rbr1-2* mutant plants.. RBR-dependent reduction or enlargement of the RM size and activation of the differentiation program align with previous observations (Hamid et al. 2025, Perilli et al. 2013). The conditional expression of RBR for 14 hours in 3-day-old seedlings was reported not to affect cell division but to promote the accumulation of the AUXIN RESPONSE FACTOR 19 (ARF19) in the root meristem, a transcription factor essential for cell differentiation and elongation (Perilli et al., 2013). However, in the same RBR-GFP line used in this study, expressing RBR via its own promoter, both the frequency of S-phase cells and the average cell size were decreased and increased, respectively, in the RM of older seedlings (5-10 DAG) (Hamid et al. 2025). Therefore, the prolonged effect of ectopic RBR may be necessary to consistently alter root growth by coordinating cell size and division.

In the epidermis of embryonic and post-embryonic cotyledons and true leaves, stomatal meristemoid cells undergo several asymmetric divisions before shifting to symmetric division and differentiating into guard cells. These small meristemoids decrease in size until they reach a threshold for symmetric division. We observed over-proliferation of stomatal meristemoids in *rbr1-2* mutant cotyledons and true leaves, forming clusters of small cells compared to the WT control. These meristemoid cells were smaller than those in WT, indicating more asymmetric divisions when RBR levels were reduced. Conversely, ectopic RBR resulted in larger meristemoids, indicating fewer asymmetric divisions with decreased meristemoid and guard cell numbers. Our data support the conclusion that the RBR level determines the size at which meristemoids stop ACD and start differentiation via SCD. Recent studies suggest that the stomatal lineage transition depends on a nuclear factor responsive to DNA content (Gong et al., 2023). Based on our data, RBR may serve as this nuclear factor, as it localises within the nucleus throughout the stomatal lineage. How does the cell cycle inhibitor RBR facilitate stomatal differentiation by promoting SCD? We observed opposing effects of elevated and reduced RBR levels on the expression of regulatory genes involved in stomatal development. Reduced RBR enhanced *SPCH* and *CYCD3;1* expression, activating and sustaining asymmetric division, while downregulating *MUTE* and *CYCD5;1*, which control the transition from ACD to SCD. Conversely, ectopic RBR expression increased *MUTE*, *CYCD5;1* and *CYCD7;1* transcripts while decreasing *CYCD3;1* mRNA levels. Thus, the upregulation of RBR contributes to stomatal meristemoid cell differentiation by regulating the expression of genes involved in the transition from asymmetric to symmetric division. Since MUTE is not a direct target of the E2F-RBR pathway (Gombos et al., 2023), the precise mechanism by which RBR controls MUTE expression remains uncertain. Nevertheless, CYCDs, involved in stomatal lineage regulation, are potential regulators of RBR through RBR kinases, whose catalytic subunits are CDKA or CDKB (Cruz-Ramirez et al., 2012; Kumar et al., 2015). RBR’s role in stomatal development may be impacted by specific phosphorylation events, leading to the formation of distinct complexes within stomatal meristemoids and guard mother cells (Simmons et al., 2019). Research shows that the CDK inhibitor SMR4 is activated by MUTE, repressing the CYCD3;1-CDK kinase to enhance G1-phase and facilitate the transition from ACD to SCD (Han et al., 2022). CYCD3;1 as an RBR-kinase component may be implicated in RBR phosphorylation, facilitating cell cycle progression. Interestingly, CYCD3;1 can bind to RBR through its LxCxE motif, whereas CYCD5;1 lacks this ability (Han et al., 2022). Consequently, the CYCD3;1 and CYCD5;1-dependent CDKs may phosphorylate distinct forms of RBR that are associated with separate protein complexes. Recent research indicates that different CDKs phosphorylate RBR at various sites, both in complex and free forms.

It has been shown that plant RBRs undergo phosphorylation at multiple sites, including 13 distinct CDK sites in Arabidopsis, similar to their animal counterparts (Pettkó-Szandtner et al., 2026). Notably, phosphorylation at the conserved 911S site is associated with multi-phosphorylated RBR forms that have lost their ability to bind and inactivate E2F transcription factors (Pettkó-Szandtner et al., 2026). In line with this, phosphorylation of RBR at this site has been shown to strongly correlates with cell proliferation (Ábrahám et al., 2011; Magyar et al., 2012; Őszi et al., 2020; Gombos et al., 2023; Pettkó-Szandtner et al., 2026). Interestingly, we observed that phosphorylation at this site also significantly correlates with RBR levels. We propose that cells begin cycling when RBR is substantially inhibited by multi-phosphorylation, as entry into the cell cycle depends on RBR’s inability to bind E2Fs. Given the relatively minor impact of different RBR levels on the development and morphogenesis of the examined transgenic plants, we suggest that higher RBR levels lead to more extensive phosphorylation, while lower RBR levels require less to enable cell division. Furthermore, since the cells in the RBR-GFP line divide less frequently but are larger, whereas the opposite is true in the *rbr1-2* line, we infer that a longer period is needed to suppress the increased RBR protein in the ectopic line, while this process happens more rapidly in the mutant compared to the WT. This may influence the duration of interphase and, consequently, cell size. Consistent with this hypothesis, a recent study has demonstrated that reduced RBR levels shorten the G1 phase in root cells (Echevarría et al., 2025). Cell size regulation in plants occurs not only during the G1 phase but also during the G2 phase, facilitated by the control of SMR CDK inhibitors through the MYB3R-SCL28-SMOS1 pathway (Nomoto et al., 2022). Therefore, two waves of CDK activities regulate cell size during the G1 and G2 phases by affecting their durations (Jones et al., 2016). These CDK activities may target RBR at both transitions, as RBR controls regulatory genes in both G1 to S and G2 to M phases (Gombos et al., 2023). Furthermore, the 911S-phosphorylated RBR forms have been identified in complexes with proteins involved in protein translation and cellular transport (Pettko-Szandtner et al., 2026). As a result, RBR, in addition to its role in cell cycle regulation, likely contributes to cell growth by controlling protein synthesis and transport, thereby influencing cell size.

We found that, in addition to cell division, the size of differentiated post-mitotic cells, such as palisade cells in true leaves and petals, was correlated with RBR levels. In a previous study, conditional reduction of RBR expression was shown to increase cell number at smaller leaf sizes due to reduced cell expansion (Borghi et al. 2010). Moreover, the effect of ectopic E2FB on leaf cell division and size was reported to be RBR-dependent (Őszi et al. 2020). Furthermore, the ARF19 transcription factor implicated in leaf expansion (Wilmoth et al., 2005) was shown to accumulate in the root in an RBR-dependent manner (Perilli et al. 2013). These observations may indicate that RBR-dependent variation in the size of dividing cells persists post-division. However, the possible effect of RBR on the elongation of post-mitotic cells cannot be ruled out. It has recently been reported that RBR levels influence hypocotyl cell elongation during the thermomorphogenetic response (Hamid et al. 2025). This was linked to corresponding changes in the expression of *YUCCA* genes associated with auxin synthesis. How RBR affects auxin synthesis-related gene expression and, consequently, cell elongation remains to be investigated.

In conclusion, this study consolidates evidence that RBR plays a fundamental role in regulating plant cell size through a level-dependent mechanism. Our results also support the hypothesis that RBR is the elusive nuclear factor responsible for regulating stomatal lineage development by balancing proliferation and differentiation by modulating key developmental transcription factors and their cyclin targets. The presented data emphasise the importance of RBR in determining cell number, cell size, and cell fate in a coordinated manner, with implications for a better understanding of plant developmental processes.

## Supplementary Data

The following supplemental materials are available.

Fig. S1. RBR regulates the area of palisade cells in petals in a dose-dependent manner.

Fig. S2. RBR-GFP exhibits widespread nuclear localization in epidermal cells.

Fig. S3. RBR regulates both the size and number of meristemoid cells in post-embryonic cotyledons.

Fig. S4. RBR regulates the size and number of meristemoid cells in embryonic cotyledons.

Fig. S5. RBR regulates guard cell numbers.

Table S1. List of primers and their sequences used for qRT-PCR analysis.

## Acknowledgements

We thank Gábor Steinbach (HUN-REN Biological Research Centre, Szeged, Hungary) for his assistance with the microscopy.

## Author contributions

Z.M., and H.B.S.R., conceived the project and designed the experiments. HBSR performed most of the experiments. M.G, F.V.-N., M.G, I.D., J.M.P.-P, and Z.M., collected materials and carried out some experiments. Z.M., H.B.S.R., and A.F., wrote and revised the manuscript with input from all the co-authors. All authors have read and approved the final manuscript.

## Data availability

All relevant data can be found within the manuscript and its online supplementary data.

